# Using Fuzzy Sets to Deal with Uncertainty in Single-species, Single-season Occupancy Models

**DOI:** 10.1101/353623

**Authors:** Malcolm L. McCallum

## Abstract

Occupancy modeling is a valuable tool for managing wildlife populations. Current occupancy models provide estimates of occurrence based on a point estimate for the species detectability and presence-absence. However, detectability can vary based on many variables ranging from weather to personnel. Therefore, I propose the use of fuzzy sets rather than point estimates for detectability and binomial presence-absence data during calculations of occupancy. Fuzzy occupancy estimates are easier to determine, more robust, and generally more informative than traditional point-based occupancy models. Consequently, managers will have better information available for comparing occupancy values among sites. Fuzzy sets are especially useful when parameters of the study violate key data standards.

## INTRODUCTION

Occupancy modeling is a valuable modeling technique in wildlife management (MacKenzie 2005). It uses arbitrarily defined spatial units or discrete naturally occurring sample units to estimate with imperfect detectability what fraction of a given area that a species occupies (MacKenzie et al. 2002). One important factor in constructing an occupancy model is determining the detectability of the species in question (Mazerolle et al. 2007). Accurately estimating this statistic is central to producing a robust occupancy model with portrays closely the actual state of occupancy in the study area (MacKenzie et al. 2003; MacKenzie 2005). The presence of a species is confirmed by detecting it on the study site. However, confirming that a species is actually absent from a site when it was not found is in most cases nearly impossible (MacKenzie et al. 2006). To deal with this problem, an array of mathematical methods is available.

If the detection probability is 1 or known to be without error, an unlikely scenario, our confidence that detection equals presence will be high. However, the more difficult an organism is to detect, the less confident we can be that sites without detection are sites without presence. In surveys for wildlife, especially endangered wildlife, it is often more important to ensure absence than presence. If the organism is absent, no special action may be needed. However, if there is any chance an exceedingly rare or difficult to detect species (e.g., Ivory Billed Woodpecker) is present, it is critical that actions be taken to avoid extinction.

There are many occupancy models and methods of calculating detectability (MacKenzie et al. 2006). Although occupancy modeling, especially with the program PRESENCE or MARK, is not difficult or hard to understand, data sets must fit a series of standards to be used in the mathematical computations. First, problems arise if you are dealing with a small number of study sites, typically 10-20 is a minimum depending on the p-value. Further, if the parameter estimates are close to 1 or 0, such as when there is a very low detection probability, psi estimates and standard error will be absurdly high. Further, all current occupancy models and their detectability estimates use and produce point estimates. A point estimator alone is often insufficient to provide a complete inference because of the importance uncertainty plays in the results (Casella and Berger 2001). The use of fuzzy sets and interval estimates in occupancy and detectability calculations allows their use when data sets are not suited and when detectability is difficult to assess.

Both interval analysis and fuzzy math are applicable to all kinds of uncertainty, without the need for subjective interpretations used in Monte Carlo approaches, and its foundation is a consistent axiomatic system that is different from that used in probability theory (Ferson et al. 1999). Fuzzy arithmetic is a conservative, nonsubjective generalization of interval analysi. Both deal well with uncertainty and require fewer data than alternative methods (Silvert 1997, 2000; Ferson et al. 1999). Whereas interval analysis uses a simple range of values, all of which are equally ranked in regard to their possibility of being true; fuzzy arithmetic rates data (*x*-axis) based on degrees of possibility (P) called membership values (*y*-axis) where *y* = 0 = lowest possibility (P_min_) and *y* = 1 = highest possibility (P_max_). All other *x*-values have decreasing membership values (i.e., possibilities, as *y* approaches zero). Fuzzy arithmetic is particularly useful where high levels of uncertainty such as ambiguity, nonspecificity, discord, and fuzziness exist (Klir and Yuan 1994). Fuzzy arithmetic was used to compare current declines with background extinction (McCallum 2007; 2015), estimate the impacts of climate change on wildlife (McCallum et al. 2009), and for risk assessment (Ferson et al. 1999).

The real advantage of fuzzy and interval-based studies is that it provides a much clearer understanding of the situation even if relatively little information is known. For example, when extinction estimates were compared with the fossil record, the actual values for extinction estimates that were calculated were very wide; however, they clearly demonstrated that current losses were not normal and compared more favorably with that previously observed during mass extinctions (McCallum 2007; McCallum 2015; Barnosky 2006; Ceballos et al. 2015). Likewise,

There are numerous mathematical approaches to occupancy modelling, all of which involve point estimates at the current time (see review in MacKenzie et al. 2006). Each has weaknesses and strengths when used with point estimates. However, with the use of fuzzy sets and intervals, they become more robust estimators.

Herein, I demonstrate using RAMAS RiscCalc (Applied Biomathematics, Setauket, NY) how fuzzy sets can improve detectability estimates used in occupancy modeling.

## A FUZZY APPROACH TO DETECTABILITY

There are many complex occupancy models for wildlife managers from which to choose (MacKenzie et al. 2006). The most basic model:

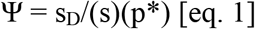

Where:

s = the number of sites surveyed

s_D_ = the number of sites where the organism was detected

p* = the probability of detecting the organism in ‘t’ surveys = 1-(1-p)^t^

If I surveyed 16 sampling sites in a preserve and detected the organism at least once in eight of those locations over three survey periods, with an estimated p = 0.8, then

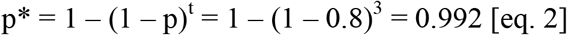
then the point estimate for occupancy at the preserve is:

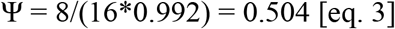

However, the true detection probability for an organism is unknown and we can safely assume that for many organisms the instantaneous detection probability on any given survey varies with many variables such as environmental factors, physiography, and habitat complexity (MacKenzie and Royle 2005). Still, we can certainly assume, barring a case of extreme luck, that the detection probability for the organism is not any less than 0.5 in the above example because that is the actual rate that we detected the organism in this study (8/16 = 0.5). If it was less detectable than 0.5, we may not expect to detect it 50% of the time. However, the surveyors may be careless or poorly skilled, and the organism might actually be perfectly detectable (p = 1) with good personnel under typical conditions. So, the detection probability could range from 0.5 – 1 with a best estimated p = 0.8. Therefore, we can set up a fuzzy set for p and calculate p*

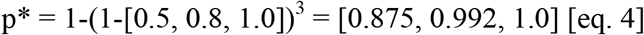

The graphical representation of p* (Fig. 1) demonstrates that detectability above 0.992 rapidly falls to zero, suggesting that values that are most representative of what is possible are < 0.992. At least, if also greater than 0.5.

The detectability estimate (p*) above is used in the following computation to calculate a fuzzy estimate for occupancy (Fig. 2):

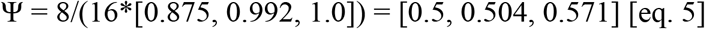

This shows that occupancy may range from 0.5 – 0. Here, Ψ = 0.504 remains the best occupancy estimate, but other rates are certainly possible. In fact, the estimates with the highest degree of possibility tend to be higher than the best estimate. This suggests that occupancy may be higher the best estimate, but it is probably not much lower than it. Notice that the base of the fuzzy estimate is equal to the interval estimate. Furthermore, there is an assurance value for ranges spanning the polygon. We do not expect the occupancy to be at or below 0.5 or at or above 0.571 because the membership values (y-axis) are zero. As occupancy approaches the best estimate (y = 1), our confidence that occupancy may fall in that range increases to 50% (y = 0.5), 70% (y = 0.7) and so on. If one was not confident in assigning a middle value for a fuzzy set, this calculation could be performed as a simple high and low to give an interval estimate instead of a fuzzy estimate. That interval estimate for occupancy would be range 0.5 – 0.571, but interval analysis will not provide the membership values on the y-axis to assess relative confidence of different values within the interval.

The above demonstration uses enough sampling sites for traditional occupancy modeling. However, if detectability is really low with a small number of sampling sites, this can be a problem with point-based occupancy models. If four sampling sites in a preserve are surveyed and the organism detected at least once in two of those locations over three survey periods, with an estimated p = 0.05, then

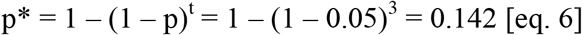

then the point estimate for occupancy at the preserve is:

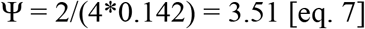

**Figure 1.**
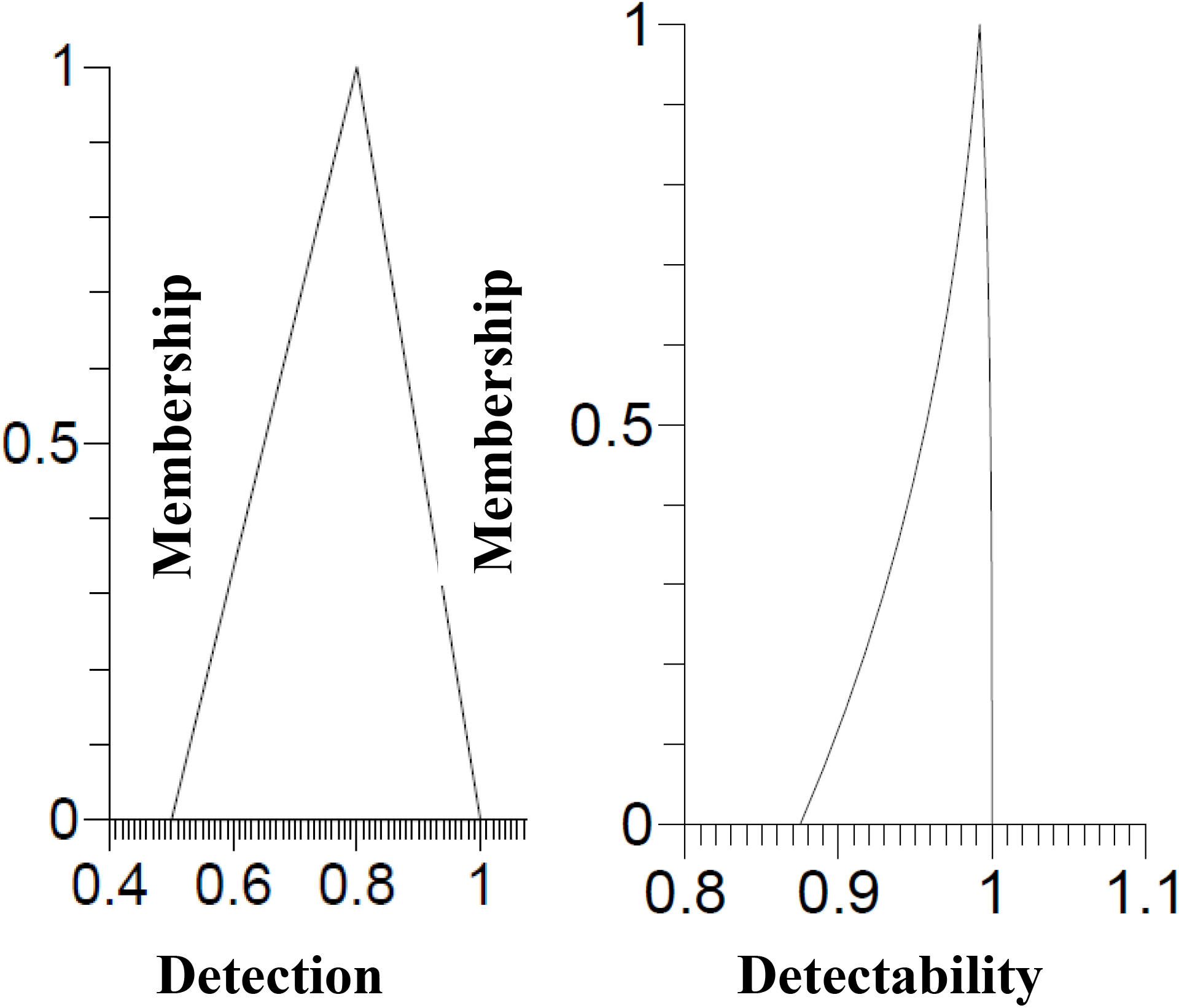
Graphical representation of p (left) and p* (right).

However, if sets are used, p* is (Fig. 3)

**Figure 2.**
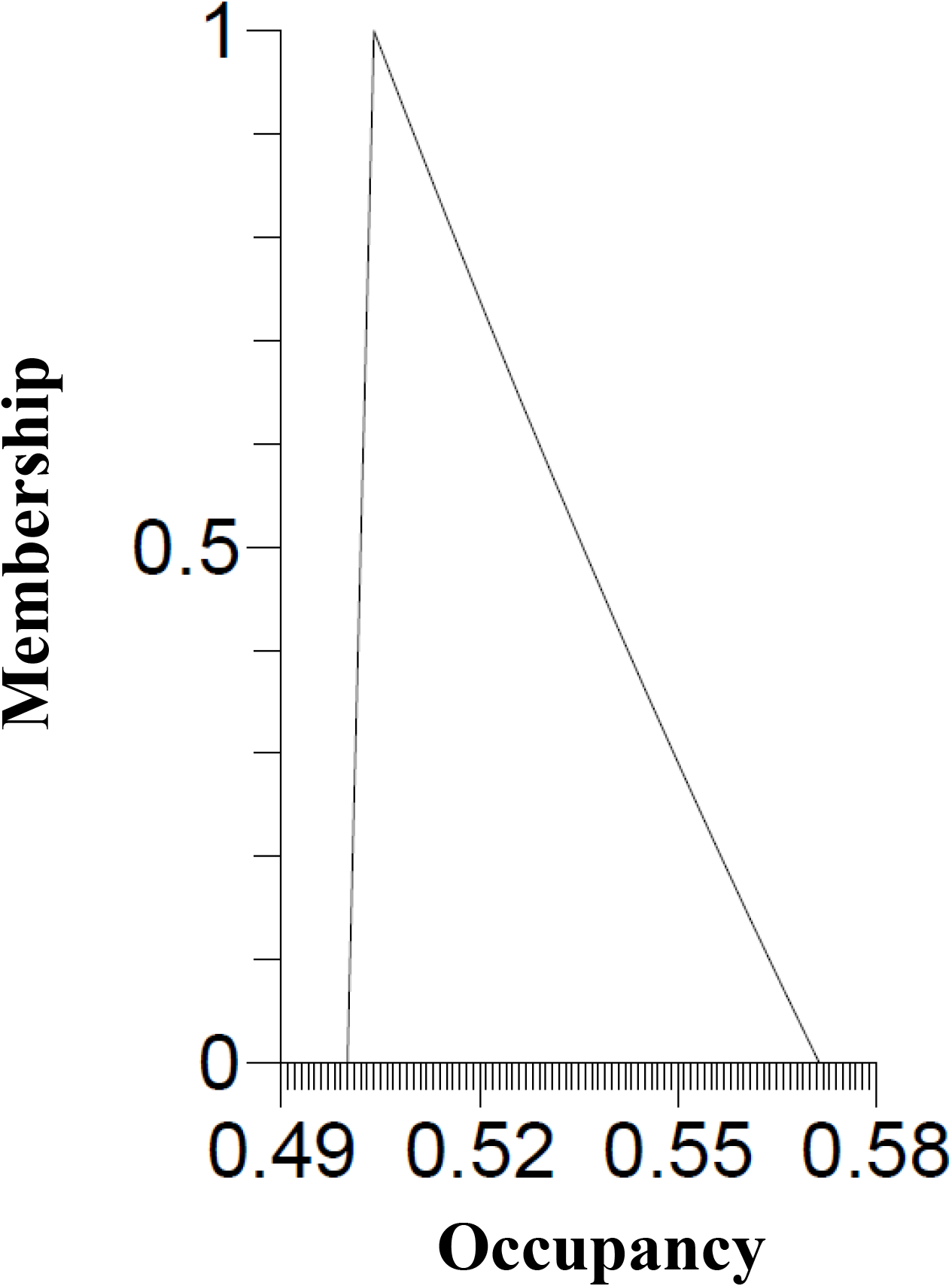
Fuzzy estimate of occupancy in a model survey site.

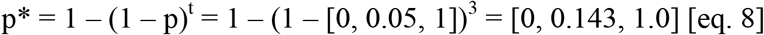

The graphical representation (Fig. 3) shows that dectability for this species won’t be much less than the best estimate; however, detectability may be substantially higher. The percent confidence is much higher for much broader intervals in this display than we saw in the earlier example because uncertainty is higher due to low numbers of sampling sites and the low detection. If an investigator does not have an informed reason for using bounds larger than zero of less than one, then it is standard practice to use a full range base on the polygon of 0 – 1. However, using zero as the lower threshold of the fuzzy or interval estimate is problematic. When p* is substituted into the occupancy formula to obtain the estimate for this preserve, we have a zero in the denominator which results in an undefined relationship.

**Figure 3.**
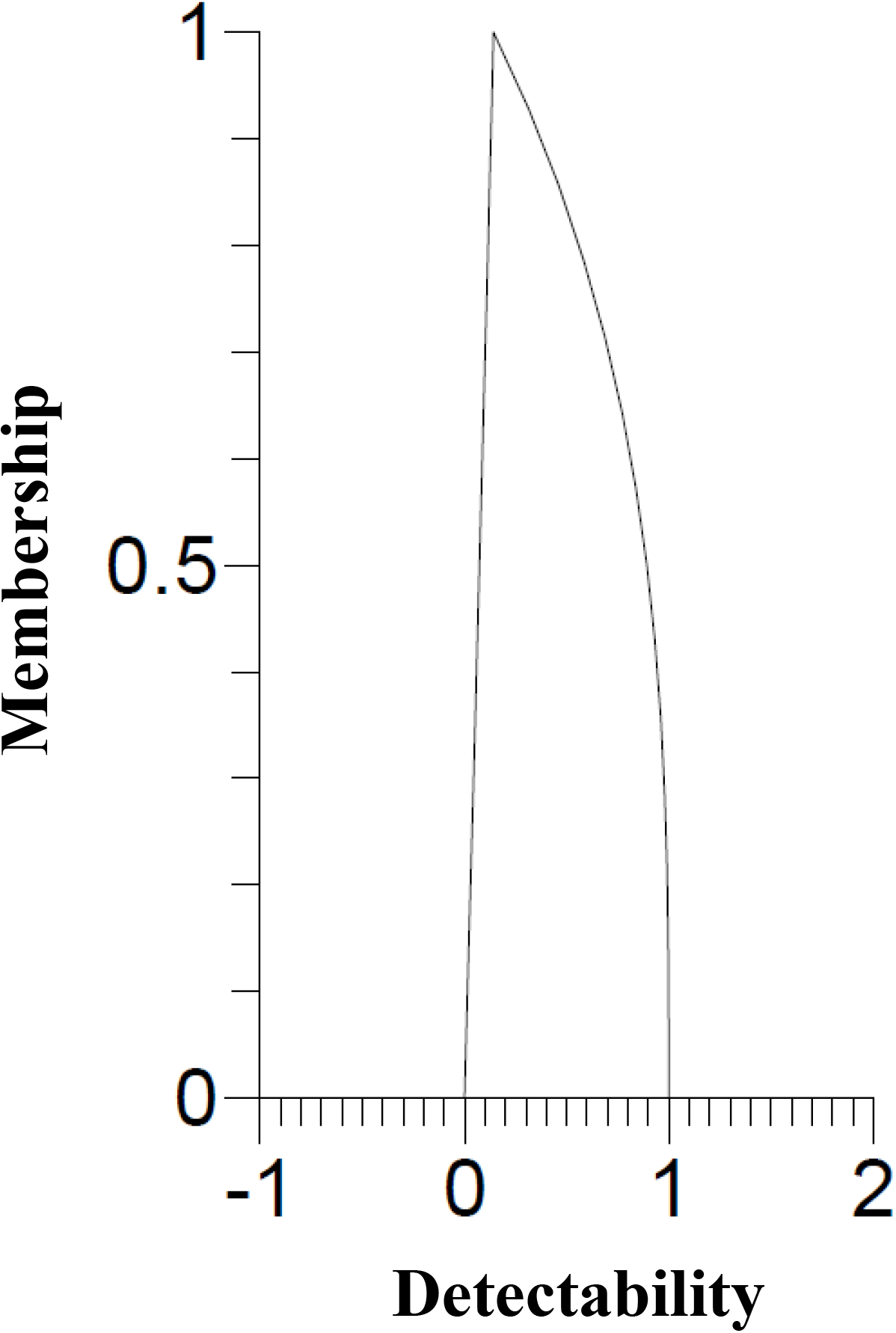
Graphical depiction of p* where detection and number of sampling sites is very low.

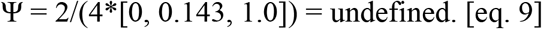

To avoid this and obtain usable information, one can substitute 0.000001 for zero.

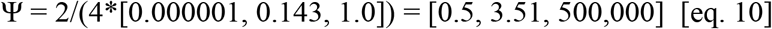

This demonstrates how such a low p* can create results that are problems for point based approaches. Essentially, there is a very high, narrow peak at 3.51, with occupancy values that slowly tail off in membership from around 10% to zero where it reaches 500,000 (Fig. 4). There is roughly a 50% assurance that occupancy is 2 – 10.

**Figure 4.**
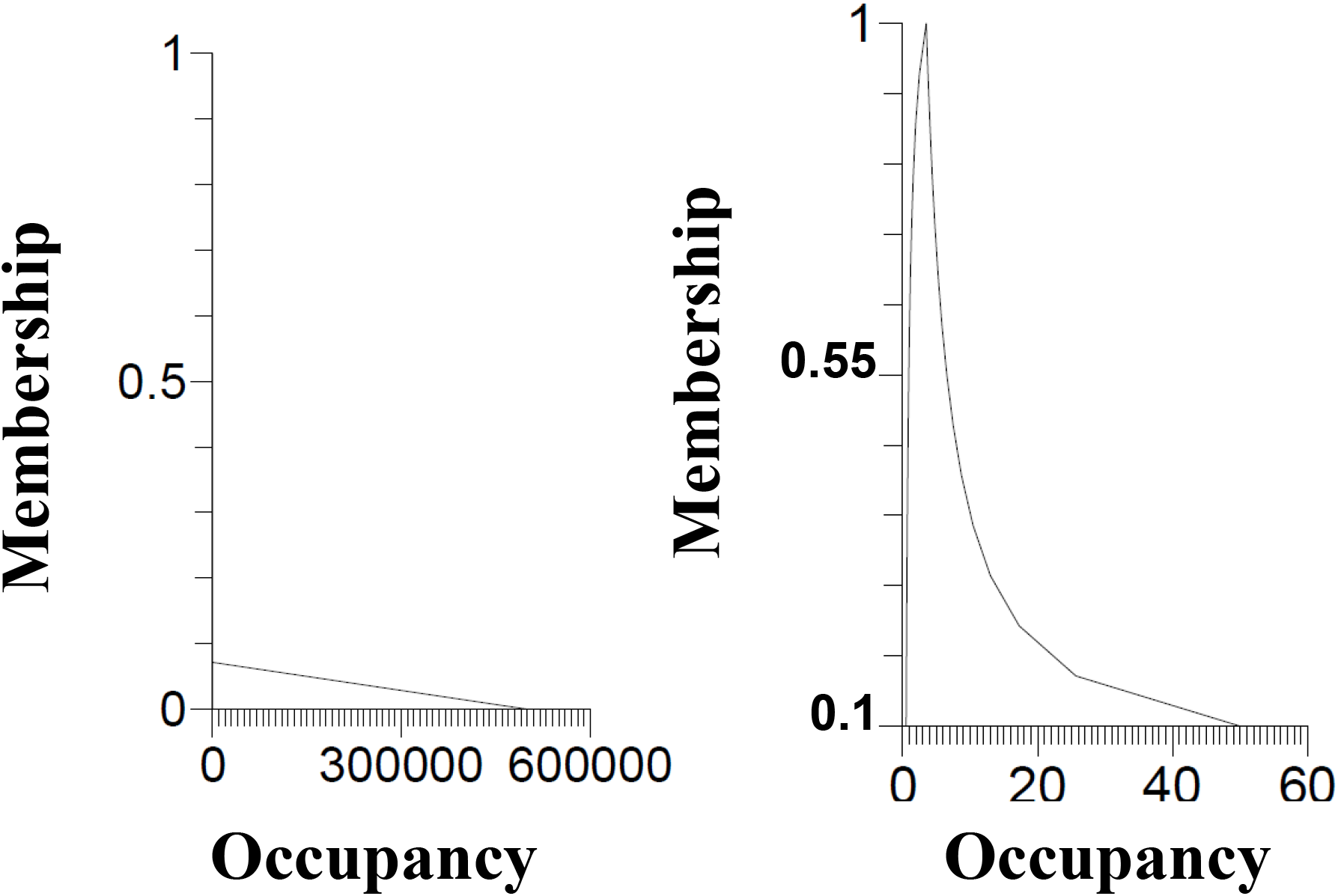
Full fuzzy occupancy estimate (left) and the portion of the fuzzy occupancy estimate above y = 0.1 (right).

This is the value of using fuzzy sets over point estimates. We know that anytime we calculate detection or occupancy that the number we generate is accompanied by a certain degree of confidence, error, or uncertainty. When we apply probability theory to occupancy, we are largely concerned with the accuracy of a single number, or what is the most probable occupancy. However, in situations where we are assessing risk of extirpation, extinction or impairment, it really is more vital to ensure absence than presence. Fuzzy and interval analyses do this. Anything outside of the bounds of the interval is generally not possible. There is much more information revealed by a fuzzy distribution than in a point estimate with error bars. As such, we can be more confident in our absence data, ensuring that actions taken do not encumber populations that were present but went undetected.

## REPLACING BINOMIAL PRESENCE-ABSENCE WITH FUZZY SETS

In the previous section, we used the binomial yes-no results from surveys in the field for one location with 16 sampling sites and another with only four sampling sites. Organisms that are difficult to detect frequently go unreported in surveys. Fuzzy sets can be used to deal with this problem. If our study site has 16 sampling locations sufficiently near to known populations we can assess the confidence in our detection or non-detection very easily. To do this, we use a sliding scale where x = 0 = did not detect, and x = 1 = did detect. If we detected the organism, then we use the fuzzy set x = [1]. However, if we did not detect the organism, we use a fuzzy set bound by zero and one [0, ?, 1] to represent the extremes that are possible, then insert an estimate on a scale of 0 to 1 representing your confidence the non-detection is accurate. How one assesses confidence may depend on the species and situation in which they are operating. For example, if I am in a parking lot and I survey it for turtles and find none, I am undoubtedly pretty confident due to the simplicity of the habitat, the type of habitat, its location relative to occupied habitat, and my ability to survey the habitat, that the non-detection is accurate. In this situation, I am very confident, so I set up a fuzzy number [0, 0.05, 1] because I am at least 95% confident that my result is accurate. However, if I were in a very complex habitat, under time constraints with a crew that was very inexperienced, I may not be very confident that nondetection is accurate. In such a situation, I account for this by estimating my confidence with a fuzzy number [0, 0.5, 1] meaning that I am only 50% confident that the non-detection is accurate, and that the species does not occur here. At first, this might seem very subjective, but converting cognitive estimates into a quantitative scheme in this way can be very accurate and informative within the study. In this study site, we had 16 sampling sites. There are various reasons you might choose to use a fuzzy estimate instead a 1 or a 0 in the calculations (Table 1.). However, to remain objective, one should develop a rubric that delineates values based on clear-cut easy to interpret standards. If we were using binomial data, our results for this inventory would 9/16 sampling sites where the animal was detected at least once during the three visits. However, in looking at the notes, we can see that some of these detections and non-detections have problems. Do we record the animal in the tree? Some investigators would be tempted to do so, but this violates the experimental design. However, ignoring its presence in the calculations is counter to the purpose of the study when we can see clearly that the animal is present and undoubtedly uses sampling site. There is even a detection that we have in our notes but makes no sense. Either the animal was well outside its normal habitat, it was misidentified, or there was an error in data recording. With the fuzzy set, we can incorporate some sense into this model to account for our certainty in my results. The fuzzy results for this inventory would give us eight sites where detection was thought accurate. Then, we have one site where the detection is baffling, and another seven sites where non-detection varies in confidence due to experience or difficulty in surveying.

**Table 1.** Fuzzy representation of detection estimates at sixteen hypothetical sampling sites for a single species within a single study site.

| Sampling Site | Fuzzy Detection Estimate | Hypothetical Rationale |
| --- | --- | --- |
| 1 | [1] | Detected |
| 2 | [1] | Detected |
| 3 | [0, 0.5, 1] | A surveyor is sure he saw the animal rapidly dive into the creek. |
| 4 | [1] | Detected |
| 5 | [0, 0.8, 1] | The animals are present in the pond just outside the sampling site. Even though we did not detect them in this site, the habitat is perfect and they are present so nearby that it is difficult to believe they do not enter this area. |
| 6 | [1] | Detected |
| 7 | [0, 0.01, 1] | The habitat is very marginal at best, and we saw no signs of the animal |
| 8 | [1] | Detected |
| 9 | [1] | Detected |
| 10 | [0, 0.9, 1] | The animal was in a tree, five feet outside our search radius. |
| 11 | [0, 0.2, 1] | We detected the animal the first time, but there are no signs of any here now, and frankly, the habitat is just not right for this species. |
| 12 | [1] | Detected |
| 13 | [1] | Detected |
| 14 | [0, 0.05, 1] | Did not find them and this just isn't a good place for them. |
| 15 | [0, 0.05, 1] | See #14 |
| 16 | [0, 0.05, 1] | See #14 |

Using the p* from [eq. 4] to calculate occupancy and then using the fuzzy set estimates for detection in the numerator

**Figure 5.**
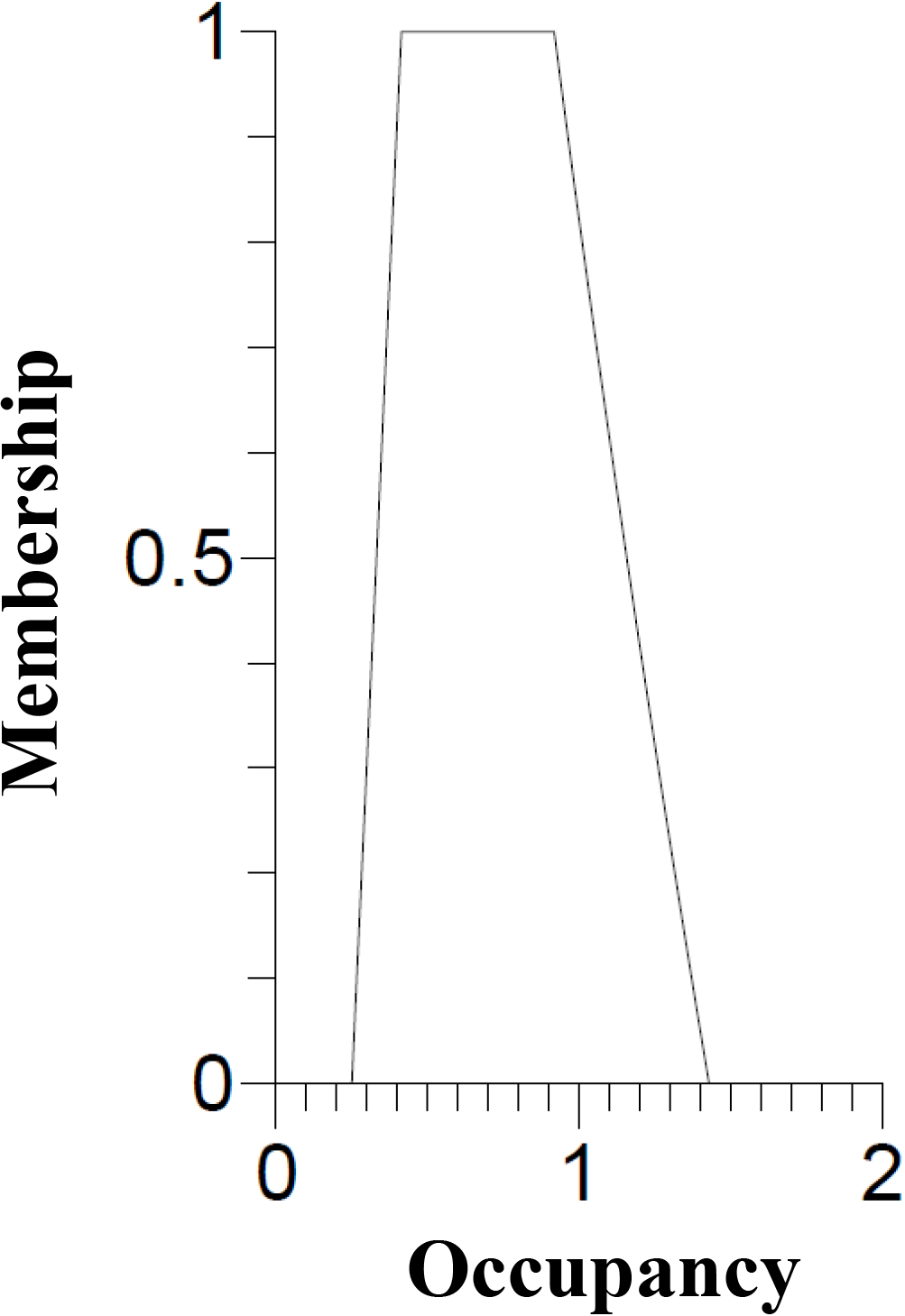
Fuzzy occupancy estimate for study site.

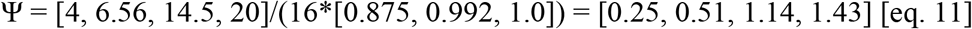

There is much more information available in the fuzzy occupancy estimate [eq. 11, Fig 5] than in a point estimate [eq. 3]. The fuzzy estimate provides that total occupancy could fall from 0.25 to 1.43, but the most credible interval estimate is 0.51 – 1.14. Any values where 1.43 < x < 0.25 are not possible. It also shows that there are more possible values > 1.14 than possible values less than 0.51. This tells us much about the status of the species that we have surveyed compared to a simple point estimate for occupancy, even if standard deviation, standard error, or confidence intervals are determined. In fact, the fuzzy estimate [0.25, 0.51, 1.14, 1.43] suggests occupancy is actually much lower than previously predicted occupancy ([0.5, 0.504, 0.571]) using a fuzzy detectability estimate and binomial presence-absence data in [eq. 5] (Fig. 2).

In addition to a more informative estimate for occupancy, fuzzy occupancy calculations are much easier to accomplish and more flexible with number of sampling sites and low detectability. In fact, the robustness of fuzzy estimates to deal with uncertainty can also encompass all kinds of variability among sampling sites. They are also much easier to interpret and communicate to others.

There are many different ways to calculate detectability and occupancy, in this study, I used a very simplistic model. However, incorporation of fuzzy sets or interval sets into more complex models is as simple as shown. However, besides fuzzifying the variables as I have done, one can also make coefficients in the functions fuzzy. The approach is the same, but this is probably not needed except under very unique and unusual circumstances.

